# Identification of the interaction of 2-aminosteroids with G-quadruplex in BCR/ABL gene promoter: An emerging potential target for treatment of human chronic myelogenous leukemia

**DOI:** 10.1101/310557

**Authors:** Qun He, Huihui Yang

## Abstract

Our preliminary studies have verified that the small molecules aminosteroid could inhibit the mRNA expression of bcr/abl fusion gene in CML (Chronic Myelogenous Leukemia) cells, which may be effective in treating CML and that may have dramatically different mechanism underlying the effects by tyrosine kinase inhibitors (TKI) binding to the BCR/ABL protein. Therefore, the exact mechanism of how the aminosteroid inhibited the CML growth should be clarified and we pay our attention to the promoter domain of BCR/ABL gene to see if any interaction between aminosteroid and the promoter. First, it should be verified if G-quadruplex could be formed in BCR/ABL promoter region. Secondly, it is highly desirable to verify if aminoteroid could be interacted with the G-quadruplex structure. Here, we reported a novel therapeutic strategy based on the targeting of the G-quadruplex which formed in the promoter regions of BCR/ABL gene by aminosteroid compounds KH and BH, verified by using bioinformatics and computer simulation, UV-Vis absorption spectra, circular dichrosism(CD), fluorescence absorption spectra, fluorescence emission lifetime expenditure experiments and polymerase chain stop assays. It showed that G-quadruplex structures can be folded in BCR/ABL promoter regions and aminosteroid inhibits the mRNA expression of BCR/ABL fusion gene by stabilizing the structure of G-quadruplex, and then inhibiting the DNA replication and transcription. This demonstrates the theory that is the so-called “chemical gene therapy” by aminosteroid in the interaction with G-quadruplex is an emerging therapeutic protocol in treatment of chronic myeloid leukemia.

## 1. Introduction

Chronic myeloid leukemia (CML) is characterized by the chromosomal abnormality arising from a translocation between the long arms of chromosome 9 and chromosome 22 and generates the BCR/ABL fusion gene[1]. The 5′-sequences of the BCR gene including its promoter and coding sequences are fused to the 3′-sequences of the ABL gene in a head-to-tail fashion [2, 3]. Therefore, a chimeric BCR/ABL mRNA transcribed from this fusion gene is initiated on the BCR gene promoter [4, 5]. The BCR/ABL fusion protein produced from the chimeric gene is strongly associated with Ph positive leukemia and is directly or indirectly causative for this blood malignancy [6] because the protein exhibited an abnormal specific tyrosine protein kinase activity from the ABL moiety of the protein gene [7]. For the BCR/ABL fusion gene, all promoter activity localizes to 1 kb area in the 5’ -exon 1 of BCR coding sequences. This region contained six consensus binding sequences for transcription factor SP1 and two CCAAT boxes but no TATA-like sequences and the overall GC contents in BCR promoter region is 78%, of which the bases are from -735 bp to -1 bp relative to the known translation start site [8]. It was reported that the rich GC sequences in promoter rigion were easy to form a specific structure known as non-canonical four-stranded G-quadruplexe (G4) [9]. G-quadruplex structure formed from four guanine bases stabilized by Hoogsteen hydrogen bonding is one of the non-canonical secondary nucleic acid structures and is able to be further stabilized by the metal cations, which is widely existed in telomere promoters and some key regulatory regions of human genome [10, 11]. And also, the G4 plays a vital role in a number of biological processes in vivo, including maintaining chromosome stability, DNA replication, transcription, genomic maintenance, RNA translational regulation, pre-mRNA splicing and gene expression[12, 13]. Recently, G-quadruplexe has been viewed as an emerging therapeutic target due to its specific promoter location and its selective oncogene promoter inhibition [14, 15]. It was reported that the transcription of targeted gene in promoter regions can be effectively hindered by stabilizing the G-quadruplex structure with small molecule binding [16]. Small molecules as G-quadruplex probes have also been paid significant attention since accumulating evidence has been linked to G-quadruplex structure in many biological processes in vivo [17].

Our previous work have proved that a kind of aminosteroids could inhibit the BCR/ABL fusion gene mRNA expression in CML cells and then inhibit the leukemia cell growth [18, 19, 20]. Thereafter, it is desired to figure out the mechanism underlying the effects of the aminosteroid. This prompts us to pay our attention to the BCR promoter region. We speculated that non-canonical four-stranded structure known as G-quadruplex (G4) may be formed in BCR gene promoter region because of high rate of GC contents similar with the c-myc oncogene promoter region[21]. And here our study showed that G-quadruplexe can be formed in BCR/ABL fusion gene promoter region verified by computer simulation, UV, CD and PCR stop assay. The interaction of aminosteroid with G-quadruplex formed in BCR/ABL promoter region was also verified by the above methods. The small molecule aminosteroid may become a new emerging therapeutic tool to target the BCR/ABL fusion gene in treatment of chronic myeloid leukemia.

## 2. Materials and Methods

### 2.1 Computational Prediction

The nucleotide sequences of BCR/ABL, Kras, c-kit, c-myc promoters and their complementary strands were obtained from NCBI (http://www.ncbi.nlm.nih.gov/) and EPD (http://epd.vital-it.ch/). The critical promoter sequences were put into computer program prediction for G4 structure formation and G-scores were calculated by using QGRS Mapper which is an online G-quadruplex prediction algorithm. This program was employed to analyse the G-quadruplex forming potential with the maximum length of 30 nucleotide bases |(Mapper, http://bioinformatics.ramapo.edu/QGRS/analyze.php).

### 2.2 Oligonuceotides and compounds

The oligonucleotides of critical promoter sequences shown in Table 1 were synthesized and purified by Gen Script (Nanjing, Jiangsu, China).The DNA stock solutions were prepared by dissolving oligonucleotides directly in 10mM Tris HCl at pH of 7.4 with 100mM KCl followed by annealing on PCR circler (first heated at 95°Cfor 8min and cooled down at the rate of 3°C per min to room temperature, then stored at 4°C at least 6h before use) for UV, CD and other detections. Aminosteroids (KH, BH, chemical structures in Fig.1) were synthesized in our laboratory and they were registered in CA as novel chemicals. Methylene Blue (MB) was purchased from Xiangya hospital (Changsha, China). The stock solutions of aminosteroids and MB were prepared by dissolving them in ethanol and ultra-distilled pure water respectively. All of the other chemicals were of analytical reagent grade and used without further purification. Ultra-distilled pure water prepared using a Milli-Q Gradient ultra-distilled pure water system (Millipore) was used in all of the experiments.

**Figure1.**
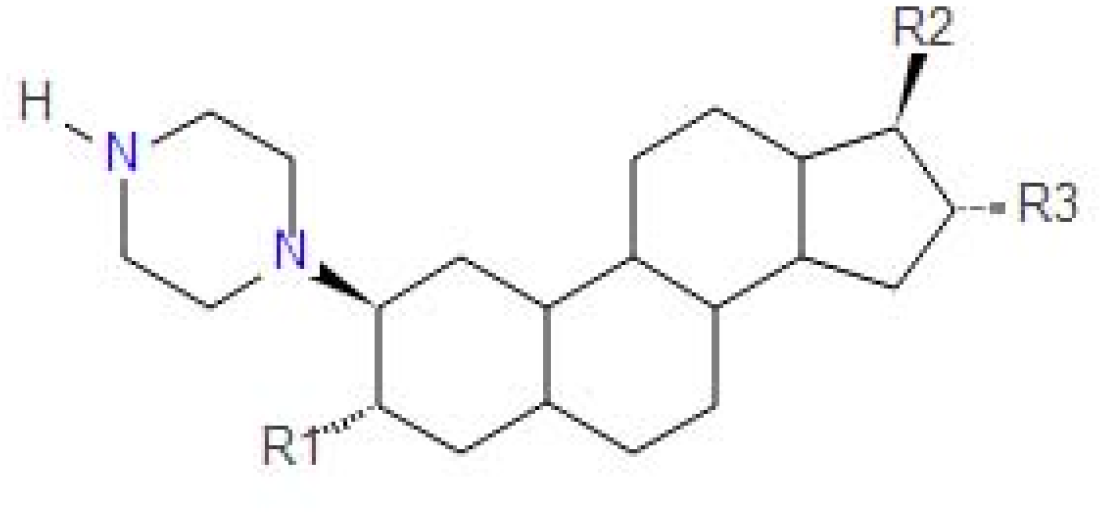
Aminosteroids (KH, BH) parental chemical structures in which R1,R2 or R3 are chemical groups such as OH, O or Br etc.

### 2.3 Circular dichroism

CD spectra were recorded on a Jasco J-815 spectrapolarimeter(JASCO, Japan) at a rate of 200 nm/min with a 0.1 cm pathlength in quartz cell at 25°C. The DNAs used in the CD experiments were at a concentration of 10uM in 10 mM Tris at pH 7.4 including 100 mM KCl. The spectra were calculated with J-815 Standard Analysis Software (Japan Spectroscopic Co. Ltd) and were recorded as ellipticity (mdeg) versus wavelength (nm). Each spectrum was recorded three times, smoothed and subtracted from the baseline.

### 2.4 UV-vis absorption spectra

The UV-vis absorption spectra (500-750 nm) detections were carried out with Varioskan Flash technique at room temperature using 96 well UV microplate (from Corning, USA). The DNAs and MB used in the UV-vis experiments were at a concentration of 10uM in 10 mM Tris at pH 7.4 including 100 mM KCl.

### 2.5 The fluorescence quenched measurements

The experiments were detected with 96 well microplate (Flat bottom, black polystyrol, from Corning, USA.) and the MB concentration was 10uM in 10 mM Tris at pH 7.4 including 100 mM KCl. All measurements were performed at room temperature and the fluorescence emission was recorded at 691nm with excitation at 666nm detected in fluorescence microplate reader.

### 2.6 Cell cultures and polymerase chain reaction stop assay

Chronic myeloid leukemia K562 cells were cultured in RPMI 1640 growth medium (Hyclone) containing L-glutamine (2 nM), antibiotics (100 U/ml penicillin, 100 mg/ml streptomycin) and 10% fetal bovine serum (Sigma). Cells were maintained in logarithmic growth phase and genomic DNA was extracted by TIANamp Gemomic DNA kit (TIANGEN biotech, China). Polymerase chain reaction stop assay was carried out with 0.5uL universe high-fidelity hot start DNA polymerase, 0.5uL dNTP mix, 0.5uL DNA template, 1uL upstream primer and 1uL downstream primer (primers used in Table 2), 12.5uL 2×unverse buffer and 2.5uL of aminosteroids, then distilled water was added to 25uL of total reaction volume, circled at 95°C 3min, 95°C15s, 60.2°C15s, 72°C 10s for 34 cycles, ended at 72°C5min and kept at 4°C. The products were separated on a 2% agarose gel prepared in TBE and the gel was exposed to autoradiograph.

## 3. Results

### 3.1 G-quadruplex can be formed in BCR/ABL promoter predicted by QGRS Mapper

BCR/ABL (HS,U07000.1), c-myc, K-ras and c-kit promoter regions and their complementary strands were obtained from NCBI and EPD, then analysed with QGRS Mapper. QGRS-mapper scans the sequence for characteristic G-rich motifs and provides a score that indicates the potential for G-quadruplex formation. The results (Table 1, Fig.2) showed that BCR/ABL promoter regions have sequences that can be potentially fold into some G-quadruplex structures referred as predicted G-quadruplex forming sequences(GQS). The G-scores in the c-myc, K-ras and c-kit promoter regions are taken as the positive controls because they were proved to form G-quadruplex structures, in which G-scores they were 42, 39 and 38 respectively. The randomised DNA oligonucleotides were taken as the negative controls which did not show any G-quadruplex forming potential with a G-score of zero because of due to the absence of four consecutive stretches of guanine bases. The GQS predicted for BCR/ABL(1018) and BCR/ABLc(78) have got a high G-score of 40 which may make its G-quadruplex as stable as that of the positive controls.

**Figure2.**
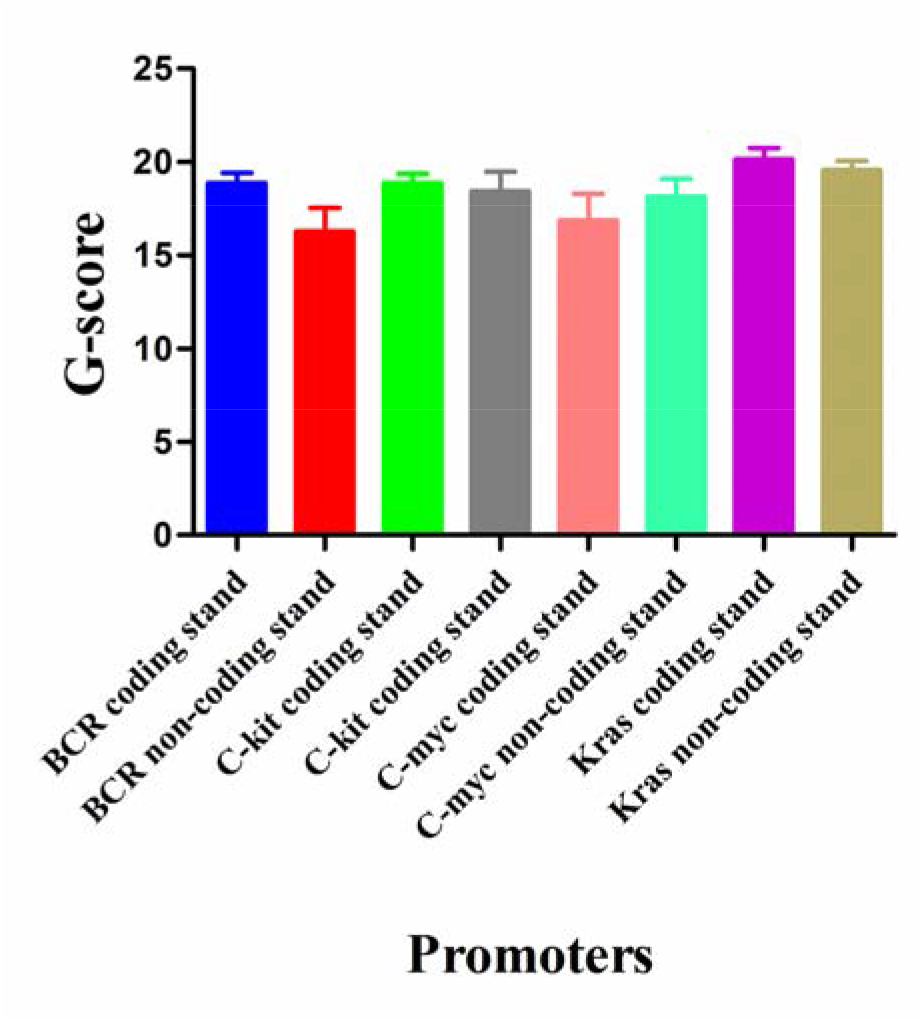
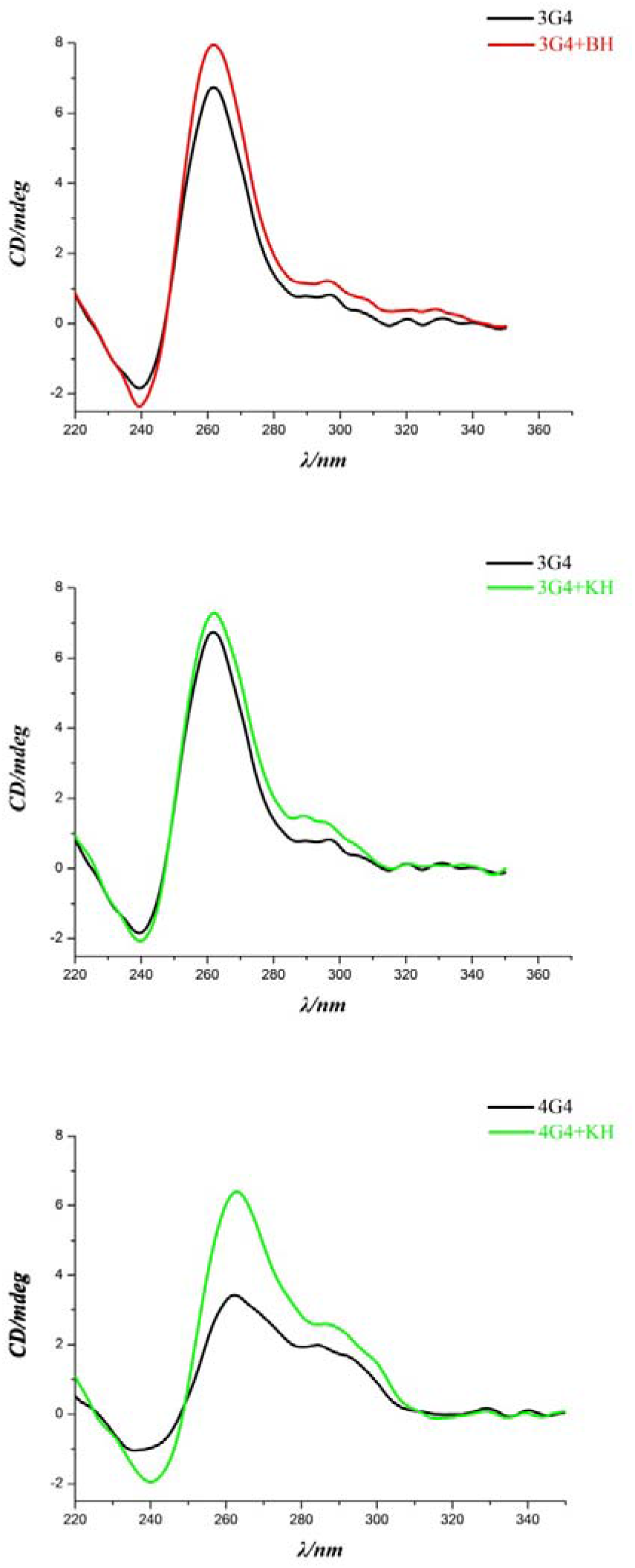
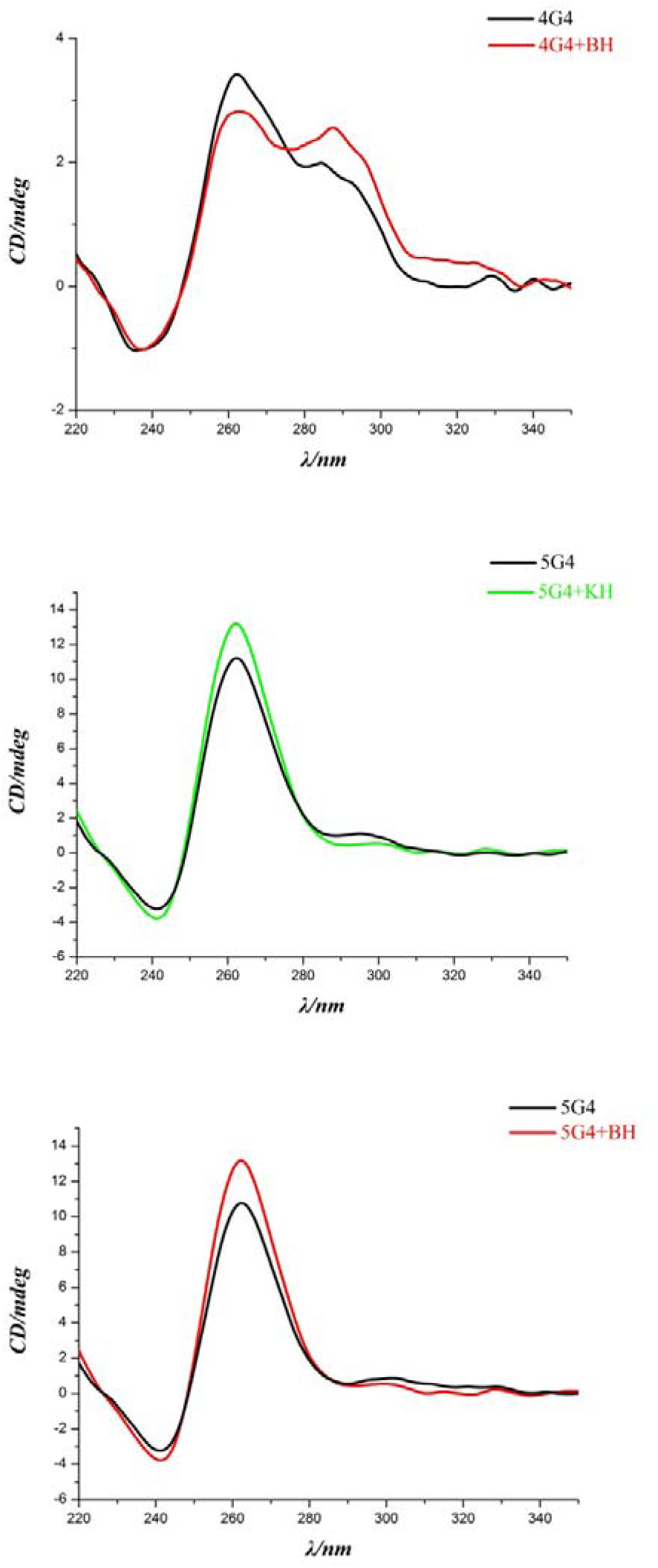
The G-scores of sequences from different promoters preticted with QGRS Mapper (The higher the score, the higher possible the G-quadruplex can be formed).

**Table1.** The G-scores of G-quadruplex forming sequences(GQS) predicted from BCR/ABL, c-kit, c-myc, K-ras, and their complementary strands. The first column showed their sequence location within their promoters and the analysis was performed with QGRS Mapper.

### 3.2 G-quadruplex structure can be formed in BCR/ABL promoter regions and the aminosteroids interacted with the structure

To verify the actual formation of G-quadruplexes predicted by QGRS Mapper, circular dichroism (CD) studies were performed on four oligonucleotides (3G4,4G4,5G4 and 8G4, Table 2).

**Table2 Oligonucleotides used in the study**

As we know, CD spectroscopy is a versatile tool to study secondary structures of nucleic acids, particularly DNA because this technique selectively distinguishes distinct conformational states of DNA depending on the salt and metal ion concentrations. CD spectra of 3G4,5G4 and 8G4 when not adding the aminosteroids showed both positive and negative CD signals at 260 and 240 nm respectively, suggesting the formation of a well-ordered parallel G-quadruplex structure. While the CD spectra of 3G4,5G4 and 8G4, when in the presence of aminosteroids, retained the both positive and negative CD signals at 260 and 240 nm respectively, but with a slight increase in CD intensity of the positive signal. CD spectrum of 4G4 showed a negative signal at 240 nm and two positive signals both at 260 nm and at 295 nm which revealed the formation of a typical hybrid G-quadruplex structure. CD data confirmed the existence of G-quadruplex structures in 3G4, 4G4, 5G4 and 8G4. Interestingly, aminosteriods could make it stable in the parallel and hybrid G-quadruplex structures (Fig.3).

**Fig.3.**
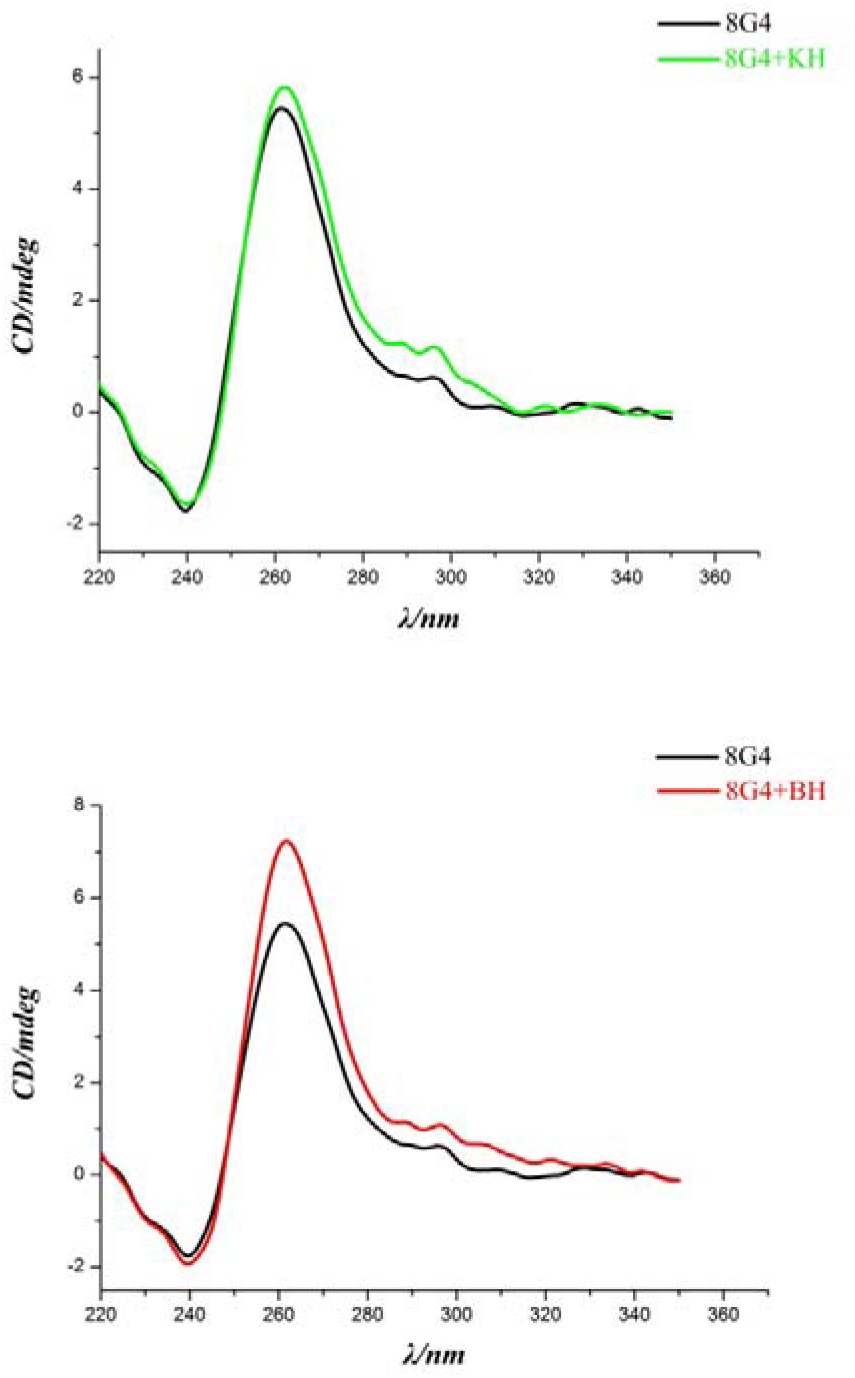
Circular dichroism (CD) spectra of different G4 in presence or not presence of aminosteroids. Concentrations: DNA(10uM), KH (10uM) and BH(10uM)

### 3.3 Methylene blue (MB) selectively probed BCR/ABL G-quadruplexes and the aminosteroids changed the shifts of MB UV or fluorescence

Methylene blue (MB) is an organic dye with a positively charged aromatic structure that belongs to the phenothiazine family and it as a G-quadruplex binding probe acquires label-free homogeneous electrochemical biosensing ability [22]. It was reported that its derivatives were taken as c-myc G-quadruplex DNA stabilizers [23], therefore we employed MB as the probe for detecting BCR/ABL promoter DNA G-quadruplexes. To investigate the feasibility of the MB probing for DNA G-quadruplex structure recognition, it was examined for its selectivity towards various DNA forms including DNA G-quadruplex sequences (3G4,4G4,5G4,8G4,7G4,6G4), single-stranded(dsDNA) and double-stranded(dsDNA), the MB probe exhibited a strong emission at 669nm when diluted in pH 7.4 Tris-HCl buffer containing 100mM KCl. In addition, MB is weakly quenching in the presence of single-stranded DNA, surprisingly, remarkable fluorescence quenching was observed after adding G-quadruplex sequences. The F/F_0_ was calculated and were as small as 0.444(dsDNA), 0.334(dsDNA), 0.170 (3G4), 0.066(4G4), 0.067(5G4), 0.218(8G4), 0.073(7G4), 0.365(6G4) respectively. The observed fluorescence quenching was due to the specific binding of the MB probe with DNA G-quadruplex structures and the fluorescence enhancement after adding aminosteroids was due to the competitive binding behavior of MB to G-quadruplexes with aminosteroid when in the presence of it. Our results suggested that MB can be used as a fluorophore to probe DNA G-quadruplex structures with an accompanying fluorescence quenching phenomena. And in this way, it was verified that the sequences predicted with QGRS Mapper in BCR/ABL promoter region could form G-quadruplex structures and the aminosteroids could bind to the G-quadruplexes that were formed (Fig.4).

**Fig.4.**
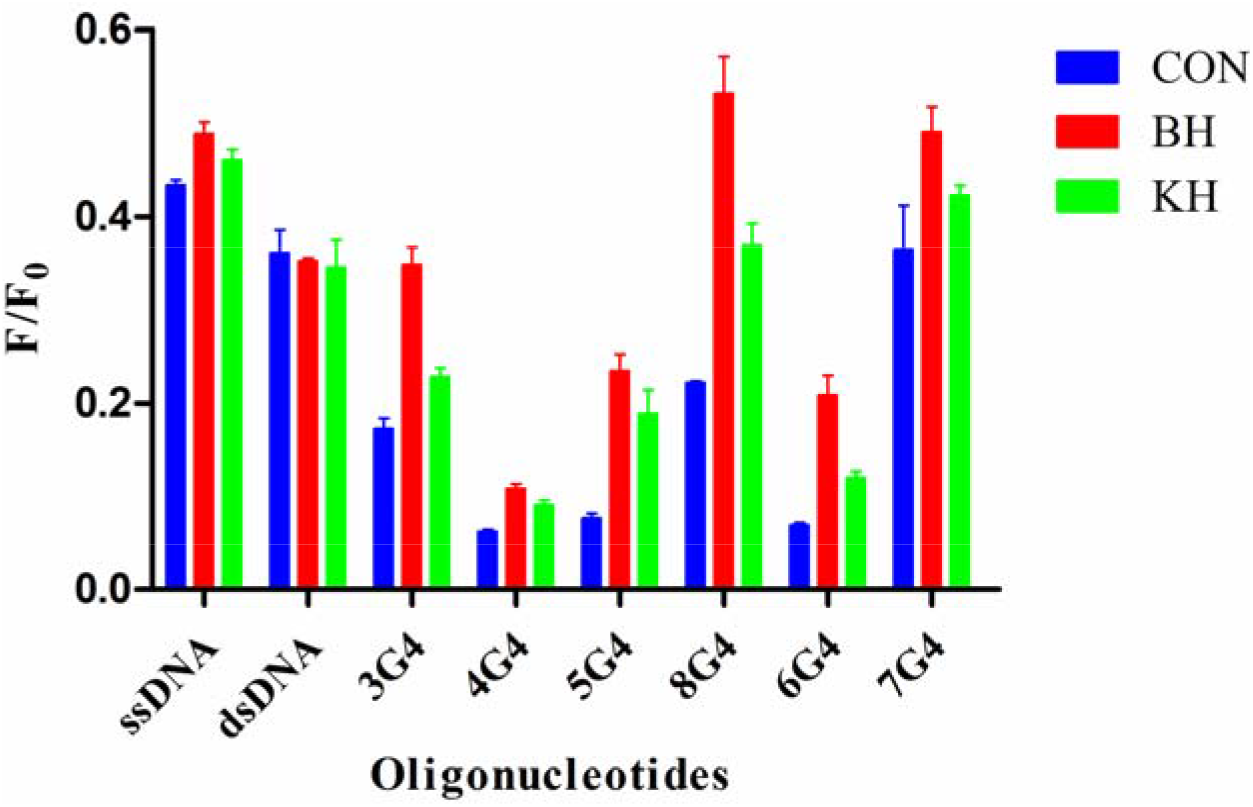
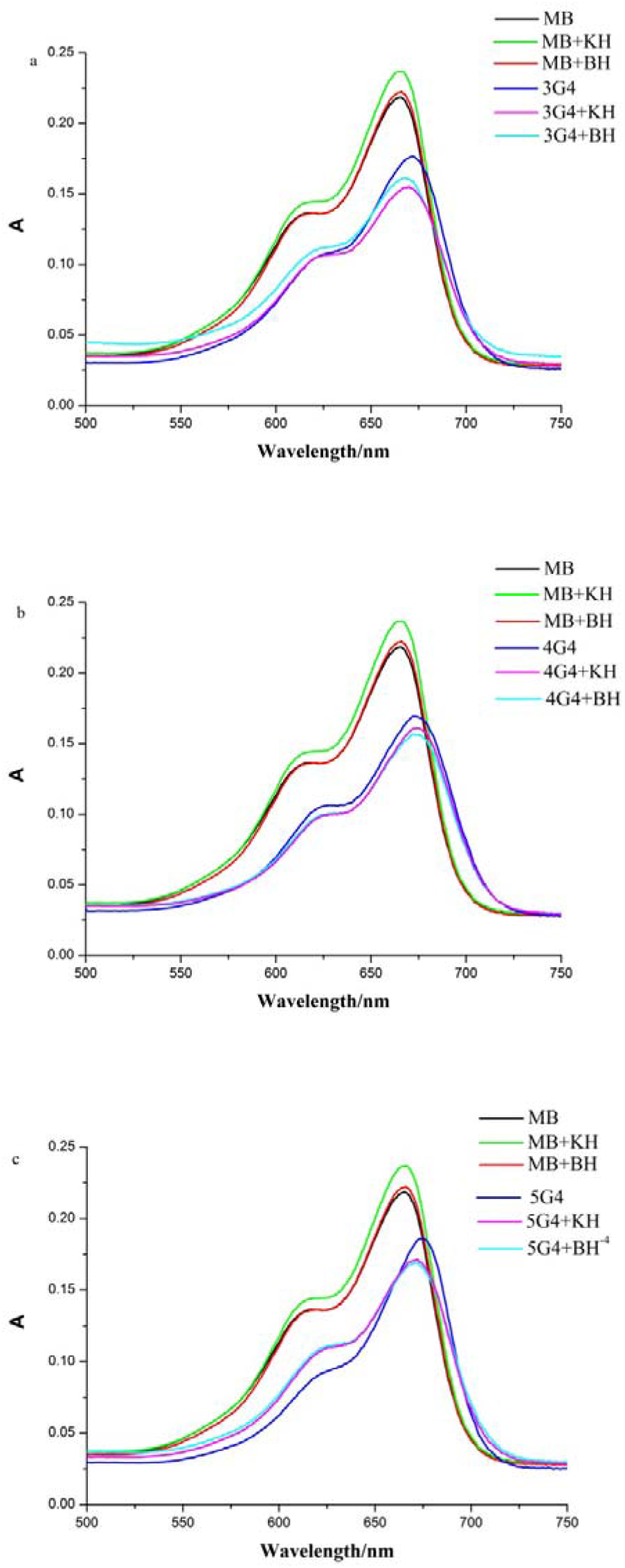
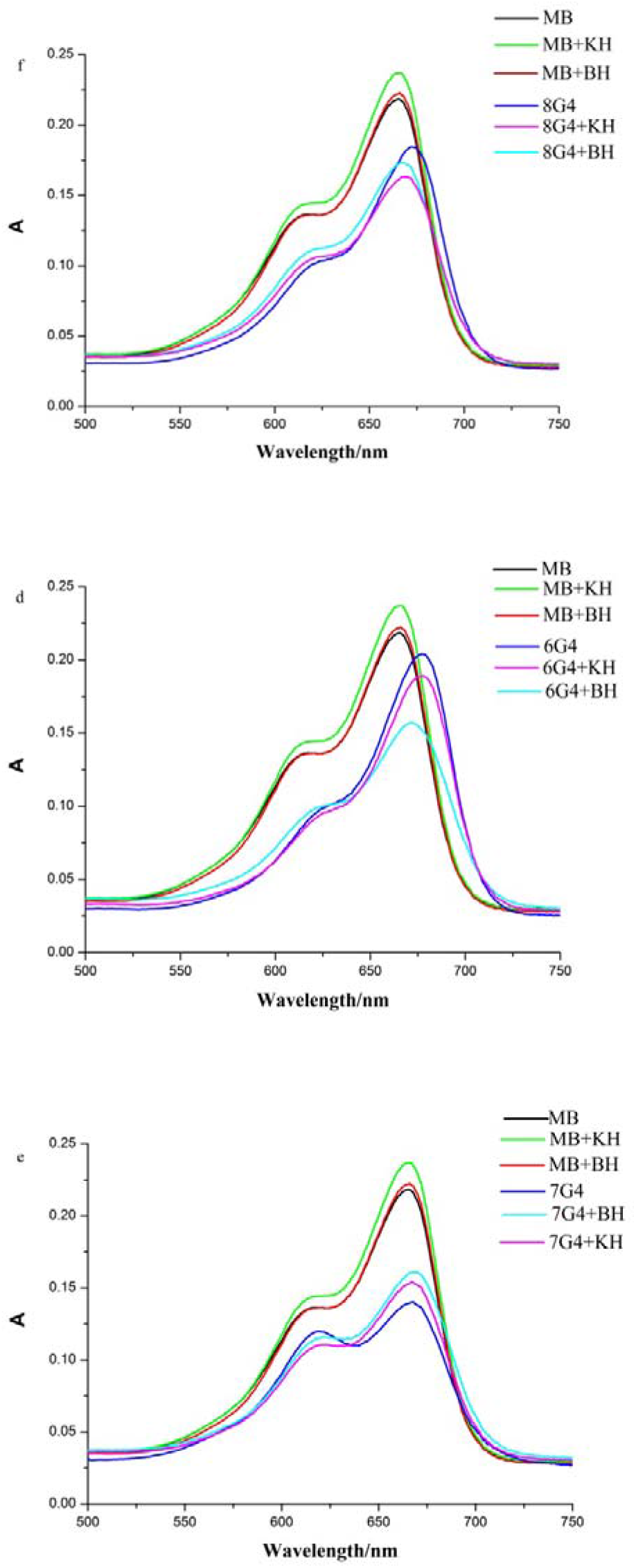
MB fluorescence quenching at 690nm for variety of DNA sequences (10uM) and the fluorescence enhancement after adding aminosteroids in 10mM Tris HCl (100mM K+, pH7.4) solution.

Furthermore, the specific probing of MB for DNA G-quadruplex structures was actually demonstrated by the UV-vis absorption spectra as shown in Fig.5, and the MB solution induced bathochromic shifts from 666nm to 672nm, 673nm, 675nm, respectively, however, blue shifts from 672nm to 667nm was observed when adding the aminosteroids to the system.

**Fig.5.**
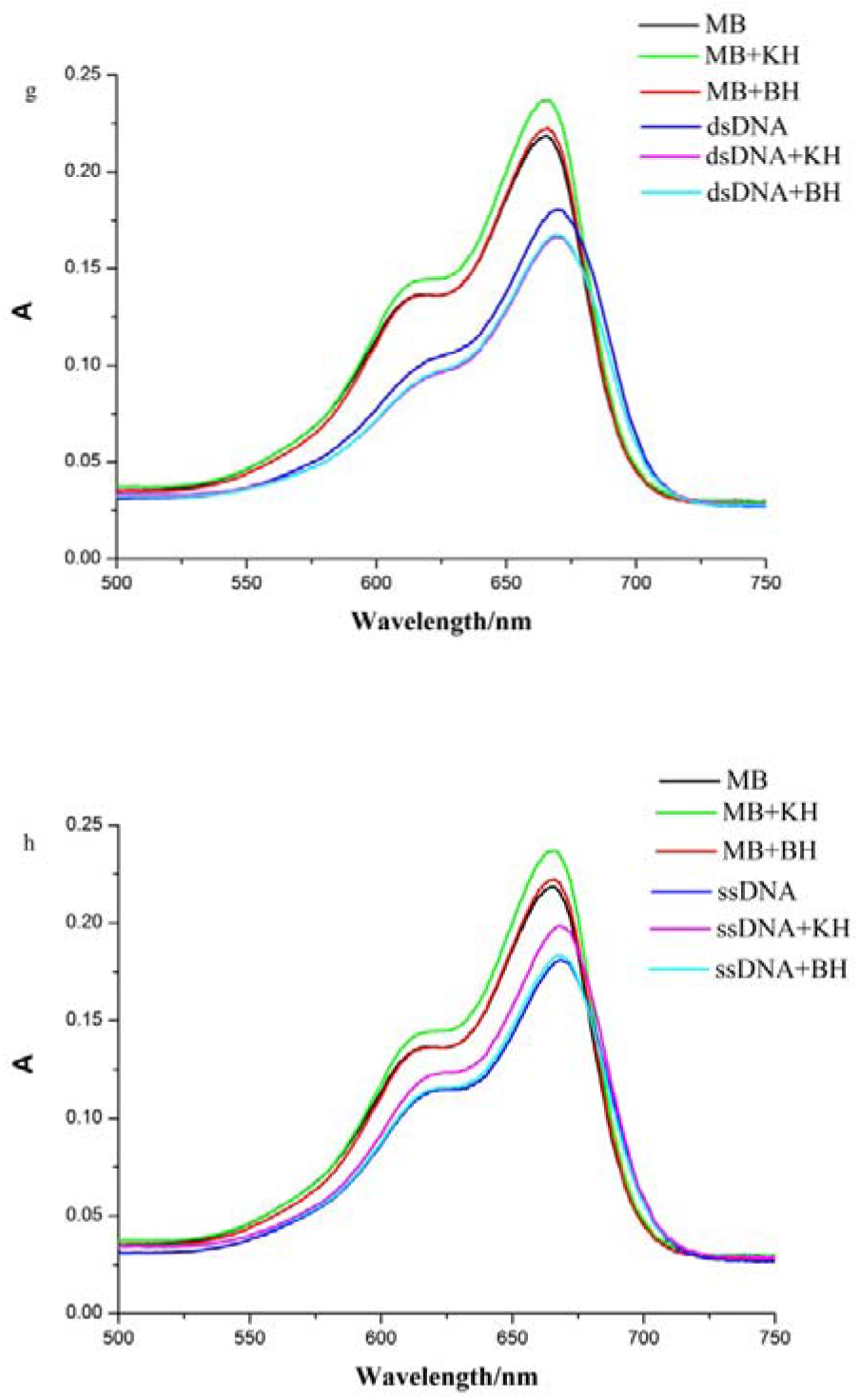
UV-vis absorption spectra of MB (10uM) with aminosteroids (100uM) and variety of DNA sequences (10uM) in 10mM Tris HCl (100mM K^+^, pH7.4) solution.

### 3.4 Aminosteroids strongly blocked primer extension in quadruplex-stabilizing status

To investigate the feasibility of the interaction of aminosteriod with BCR/ABL promoter, we examined its effectiveness using the PCR-stop assay. The aminosteriod compounds were incubated with genomic DNA extracted from K562 cells and the complementary reverse primer in the presence of Universe High-Fidelity Hot Start DNA Polymerase. Stabilization of the intramolecular G-quadruplex structure by KH and BH prevents hybridization of the complementary sequence. Universe High-Fidelity Hot Start DNA Polymerase is unable to recognize the G-quadruplex structure and DNA amplification is inhibited, which is manifested as a reduction in the 303 bp PCR product observed after agarose gel electrophoresis (Fig6). The results show that the addition of aminosteroids led to a decrease in the 303 bp PCR amplification product. Partial or total inhibition of Universe High-Fidelity Hot Start DNA Polymerase-mediated DNA amplification through stabilization of the G-quadruplex structure was observed at 100 uM of KH and BH, and especially complete inhibition by BH. By comparison, This indicates that aminosteriod are able to stabilize G-quadruplex formation in biological systems. Furthermore, this result confirms that the BH binds more strongly to the BCR/ABL G-quadruplex compared to KH, which is in agreement with our other experiments.

**Fig.6.**
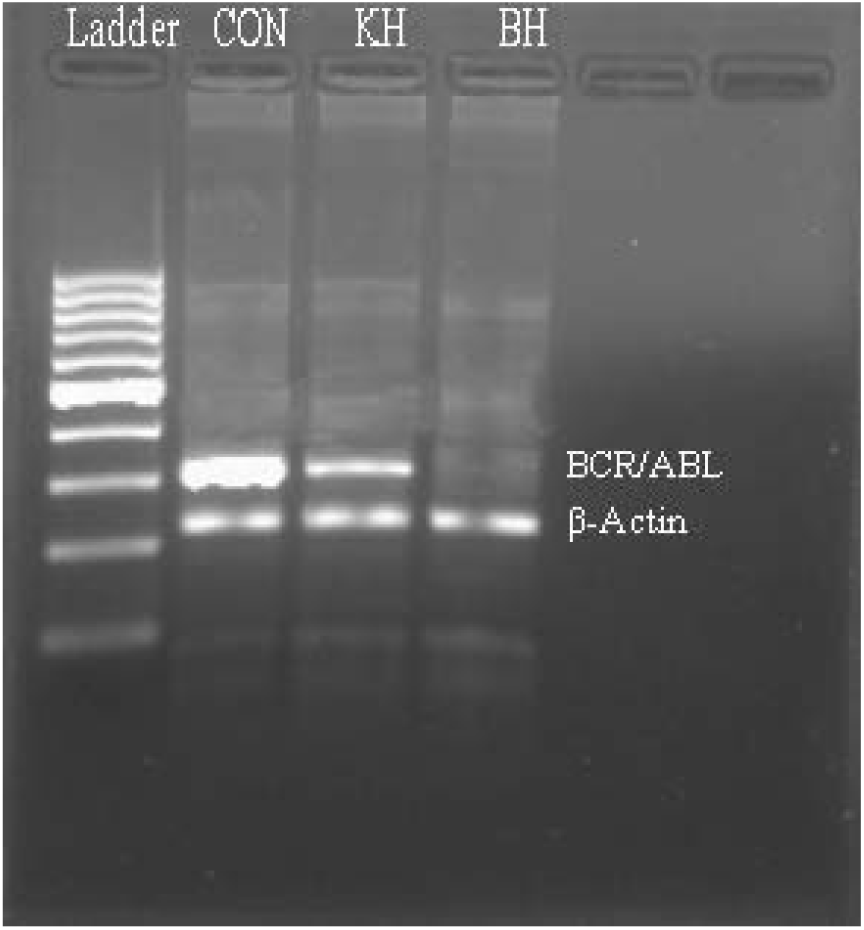
PCR stop assay for BCR/ABL and β-actin in genomic DNA of K562cell lines after no treatment and treatment with 100uM BH and KH respectively.

## 4. Discussion

At present, most researches have been committed to develop precise target treatment by chemotherapies on BCR/ABL in CML. Over the past decades, some BCR/ABL inhibitors such as tyrosine kinase inhibitors (TKIs) have been used in clinic worldwide. Though TKIs can improve the life expectancy of patients suffering from chronic myeloid leukemia (CML), patients would eventually develop resistance to TKIs therapy or would not endure the adverse effects due to secondary off-target mechanisms associated with TKIs [24].

One of resistance mechanisms was the BCR/ABL protein configuration changed resulting in the TKIs could not anymore bind to the protein. Our small molecules such as KH and BH target the BCR/ABL promoter gene, but not the protein. Therefore, even the resistant CML cells would respond to our molecules as verified by our previous papers. G-quadruplex structures have emerged as a new anti-cancer strategy due to their vital biological function in cancer cells proliferation. More and more accumulating evidences were showing that the researches emphasize on G-quadruplex detection, imaging and its interference with some small molecules in living cancer cells [25]. However, it was seldom reported in leukemia such as CML.

Prediction of potential GQS relies on the folding rule: GxNy1GxNy2GxNy3Gx [25]. In which G represents guanine residues taking part in G-tetrad formation and x is the number of successive guanines. According to the folding rule, x is always constant and should be at least two because a minimum of two quartets is required to stack on top of each other to form a G-quadruplex. The lower scoring GQS will have two G-tetrads in its G-quadruplex entity and the stability is enhanced by more G-quartets or the number of guanine residues participating in the stacks. N is representative of the other bases involved in the loops of the G-quadruplex where N can be any base including guanine. y1, y2 and y3 are the frequency of the different residues that participate in the three different loops and they can vary. Loop size can affect stability of G-quadruplex ensembles but not as much as the frequency of guanine bases. GQSs having at least three guanine tetrads and loops of equal length connecting them will be highly stable and have high G-scores such as the positive controls.

CD is a powerful experimental method to examine the G-quadruplex structures as well as ligands binding of quadruplex DNAs or RNAs. It is special by comparing their spectra with those known quadruplex DNAs or RNAs structures[26,27,28]. The CD spectrum has the positive band near 260 nm and a negative band near 240 nm that are characteristics of an parallel known G-quadruplex structure. In the other respect, if CD spectrum is characteristic of a positive band near 260 and 290nm, then a negative band near 240nm, that means an hybrid structure existed (both parallel and anti-parallel structures)[29,30,31].

According to the above CD structure identification regulation, it was first discovered that the sequences in upstream of BCR/ABL promoter region which was the major transcription initiation site could be able to form four intramolecular G-quadruplex structures under similar physiological conditions. Furthermore, the G-quadruplex structures formed included two different kinds of structure topologies, namely both parallel and hybrid as verified in the results. Then, as the results, it was verified that aminosteroids KH and BH could stabilize both the G4 parallel and hybrid structures. It is therefore suggested that the interaction of aminosteroids with G-quadruplex formed in BCR/ABL promoter region may be a new potential therapeutic protocol in treatment of chronic myeloid leukemia.

Nowadays, people try to treat cancer such as leukemia by using gene therapy [32]. Generally, gene therapy needs a vector which is usually a virus to conduct a gene into cells, by which method, it could be called “biological gene therapy”. Our protocol was to use a small molecule which could directly interfere with gene such as BCR/ABL promoter gene, in this way to do so-called “chemical gene therapy”. In this method, we hope that the manipulation of targeted gene would be more simple, no vector needed and more applicable in future clinical translation. Thereafter, we found a potential way to make aminosteroids to be a novel protocol to treat CML in the future.

## Contributions

He Q was in charge of paper design, experiment direction, data analysis and article writing. Yang HH was in charge of experiment doing and data collection.

## Conflict of interest

All authors declare no conflict of interest to other people or other journals.

## Acknowledgements

This work was supported by the Grants from Natural Science Foundation of China (39970316, 30470638 and 81070483)

